# Validation of a novel milk progesterone-based tool to monitor luteolysis in dairy cows. Timing of the alerts and robustness against missing values

**DOI:** 10.1101/526095

**Authors:** Ines Adriaens, Olivier Martin, Wouter Saeys, Bart De Ketelaere, Nicolas C. Friggens, Ben Aernouts

## Abstract

Automated monitoring of fertility in dairy cows using milk progesterone is based on the accurate and timely identification of luteolysis. In this way, well-adapted insemination advice can be provided to the farmer to further optimize the fertility management. To properly evaluate and compare the performance of new and existing data-processing algorithms, a test dataset of progesterone time-series that fully covers the desired variability in progesterone profiles is needed. Further, the data should be measured with a high frequency to allow rapid onset events, such as luteolysis, to be precisely determined. Collecting this type of data would require a lot of time, effort and budget. In the absence of such data, an alternative was developed using simulated progesterone profiles for multiple cows and lactations, in which the different fertility statuses were represented. To these, relevant variability in terms of cycle characteristics and measurement error was added, resulting in a large cost-efficient dataset of well-controlled but highly variable and farm-representative profiles. Besides the progesterone profiles, information on (the timing of) luteolysis was extracted from the modelling approach and used as a reference for the evaluation and comparison of the algorithms. In this study, two progesterone monitoring tools were compared: a multiprocess Kalman filter combined with a fixed threshold on the smoothed progesterone values to detect luteolysis, and a progesterone monitoring algorithm using synergistic control ‘PMASC’, which uses a mathematical model based on the luteal dynamics and a statistical control chart to detect luteolysis. The timing of the alerts and the robustness against missing values of both algorithms were investigated using two different sampling schemes: one sample per cow every eight hours versus one sample per day. The alerts for luteolysis of the PMASC algorithm were on average 20 hours earlier compared to the ones of the multiprocess Kalman filter, and their timing was less sensitive to missing values. This was shown by the fact that, when one sample per day was used, the Kalman filter gave its alerts on average 24 hours later, and the variability in timing of the alerts compared to simulated luteolysis increased with 22%. Accordingly, we postulate that implementation of the PMASC system could improve the consistency of luteolysis detection on farm and lower the analysis costs compared to the current state of the art.

**Interpretative Summary:** Validation of luteolysis monitoring tool for dairy cows.

**Adriaens:** In this study, the performance of two monitoring algorithms to detect luteolysis using milk progesterone measurements was validated on a simulated dataset of realistic milk progesterone profiles. The synergistic control-based algorithm, PMASC, was able to identify luteolysis almost simultaneously with its occurrence. It was found to be more robust against missing samples and less dependent on the absolute milk progesterone values compared to a multiprocess Kalman filter combined with a fixed threshold. This research showed that implementation of PMASC could improve progesterone-based fertility monitoring on farm.

## INTRODUCTION

Monitoring of milk progesterone (**P4**) in dairy cows allows identification of a cows’ reproduction status. Because P4 is fat-soluble and transfers from the blood into the milk, the concentration of P4 in milk is 4 to 5 times higher than in blood. High P4 concentrations, produced by an active corpus luteum (**CL**) on the ovaries are associated with the luteal phase of the cycle or pregnancy, while low P4 concentrations are known to occur during the follicular phase of the P4 cycle and in the postpartum anestrus phase after calving. Luteolysis, under influence of the uterine PGF_2α_ signal and defined as the regression of the CL, is accompanied with a steep and fast decrease in P4, seen as a drop in milk P4 from over 15 ng/mL to below 5 ng/mL in approximately 12 to 24 hours. This drop in P4 is necessary to allow for a LH surge that induces rupture of a pre-ovulatory follicle (ovulation). Estrus detection based on milk P4 dynamics therefore relies on the accurate and timely identification of luteolysis preceding ovulation. Since recently, it is possible to automatically measure milk P4 on farm, in which regular milk analyses clearly show the P4 dynamics during an estrous cycle (Adriaens et al., 2017; Bruinjé et al., 2017). The current state-of-the-art in P4-based fertility monitoring is to smooth the raw measured values with a multi-process Kalman filter (**MPKF**), after which a fixed threshold (**T**) to these smoothed values is applied to detect luteal activity and luteolysis (Friggens and Chagunda, 2005; Friggens et al., 2008). The MPKF hereby ensures that no alerts are triggered for a single low measurement. The set threshold’s value might depend on the P4 measurement technique or the calibration method, but is generally taken between 4 and 6 ng/mL (Friggens et al., 2008; Bruinjé et al., 2017). In contrast, the P4 monitoring algorithm using synergistic control (**PMASC**) enables the identification of fertility events on farm using the underlying physiological basis of the related P4 dynamics (Adriaens et al., 2017, 2018a). It employs a combination of mathematical functions to describe the development and regression of the CL and a statistical control chart for detection of luteolysis. Until now, this system was designed, optimized and evaluated on high-frequent P4 measurements in which milk P4 was analyzed *post-hoc* via ELISA-testing in the lab. This did not yet represent on-farm measured data for which the measurement error is representative, the time series are sufficiently long and in which all the variability in P4 profiles on farm is included. Before the PMASC algorithm can be used on farm, it should therefore also be validated as such.

In the ideal scenario, this validation would be performed on a large dataset representative for on-farm measurements and containing numerous milk P4 profiles with as much variability as possible, not only in fertility and profile characteristics (e.g. including follicular and luteal cysts, early and late pregnancy and embryonic losses within a lactation) but also in cycle shapes (e.g. height, baseline and slopes of each P4 cycle). The frequency of measurement should be as high as possible (e.g. once per milking) in order to be able to vary sampling schemes and test all possible scenarios. Moreover, to validate a luteolysis monitoring algorithm, the actual moment of luteolysis should ideally be known in order to use this as the ‘gold standard’. To our knowledge, and especially with respect to time of luteolysis measures, a dataset does not exist that meets all these criteria.

Alternatively, a convenient and more efficient way to obtain an appropriate dataset is through simulations, which allow generation of extensive datasets while avoiding analysis costs both in terms of measurements and time. Recently, a systemic white box model representing a virtual cow (GARUNS, Martin and Sauvant, 2010) coupled to a model describing the reproductive functioning (reproduction function model, **RFM**, Martin et al., 2018) was developed. This model allows simulation of virtual cows with diverse fertility characteristics, and provides scaled P4 profiles corresponding to the reproductive functioning of the simulated cows. The RFM outputs scaled P4 profiles, meaning that the dynamics are representative for the fertility, but need to be adjusted to represent the targeted measurement technique and substrate (milk vs. blood), from which we know the absolute values can vary. In this way, large datasets representing cows with sufficiently variable fertility characteristics, for which the number of estruses and timing of luteolyses is known, can be obtained.

In a previous validation study, PMASC was shown to be successful in correctly identifying luteolysis using milk P4 data measured on-farm with a lateral flow immunoassay (**LFIA**) (Adriaens et al., 2019). Nevertheless, as those P4 data originated from a commercial sensor system, the sampling frequency was set by the system software and the effect of the measurement frequency and timing, as well as the effect of missing values on the performance of PMASC could not be evaluated. Additionally, the timing of the alerts generated by PMASC and MPKF+T could not be verified as no reference information on the exact moment of luteolysis was available on those farms. Therefore, the objective of the current study was to compare the detection performance, robustness and consistency of alerts from PMASC and the MPKF+T method based on simulated realistic P4 profiles. Furthermore, this approach allowed for the evaluation of the effect of missing samples or a reduced sampling scheme during luteolysis. A good understanding of the performance of the P4 monitoring algorithm under variable sampling conditions and schemes is important to quantify the uncertainty in the prediction of the optimal insemination window.

## MATERIALS AND METHODS

### Simulating Progesterone Profiles

In a first step, 100 dairy cows were simulated using a systemic model for describing lifetime performance in dairy cows (GARUNS), developed by Martin et al. (Martin and Sauvant, 2010; Martin et al., 2013, 2018; Gaillard et al., 2016). The fertility characteristics of these cows were defined by a recently developed reproduction module coupled to GARUNS, the RFM, for which a schematic overview is shown in Figure A1. The general idea of the RFM is that a cow shifts continuously between 11 different fertility compartments, namely ‘prepubertal’, ‘anestrous’, ‘anovulatory’, ‘pre-ovulating’, ‘ovulating’, ‘post-ovulating’, ‘luteinizing’, ‘luteal’, ‘cystic’, ‘dysfunctional’ and ‘gestating’. The dynamics of these shifts are influenced by the lifetime performance model GARUNS, for instance by the rate of hormonal clearance and the energy balance. In turn, the RFM’ outputs ‘conception’ and ‘embryonic/fetal death’ manipulate the course of GARUNS, and thus the life of a cow, to trigger body weight changes, dry-off, initiation of a new lactation and so on. The competence stages ‘cystic’ and ‘dysfunctional’ are modifications of the original published work (Martin et al., 2018) that allow for interruption of cyclicity in absence of P4 and a P4-producing (luteal) cyst-like structure, respectively. The first can be either sudden anestrus (no activity on the ovaries) or a follicular cyst (large, fluid filled follicular structure on the ovaries) (Ranasinghe et al., 2011; Jeengar et al., 2014). The latter results in intermediate P4 concentrations during the luteal phase as described by Braw-Tal et al., (2009), Peter et al., (2009) and Rosenberg, (2010). The parameters of the model (Table A1) were chosen to obtain cows with a large range of fertility characteristics, both in terms of length of the postpartum anestrus period, number and length of the cycles, occurrence of interrupted cyclicity due to follicular and luteal cysts and the interval to successful pregnancy.

Each simulated cow had 6 to 7 lactations with different fertility features reflected in the scaled P4 profiles (example in Figure 1, red). A subset of lactations was selected based on the variability in P4 profile characteristics. For this, the profiles were successively sorted by postpartum anestrus length, number of cycles and incidence of interrupted cyclicity, after which each time the 50 most variable lactations were selected. This resulted in a dataset of 150 profiles (i.e. the consecutive dynamics over 1 lactation) containing in total 731 scaled estrous cycles. The dataset characteristics are summarized in Table 1.

**Adriaens, Table 1.**
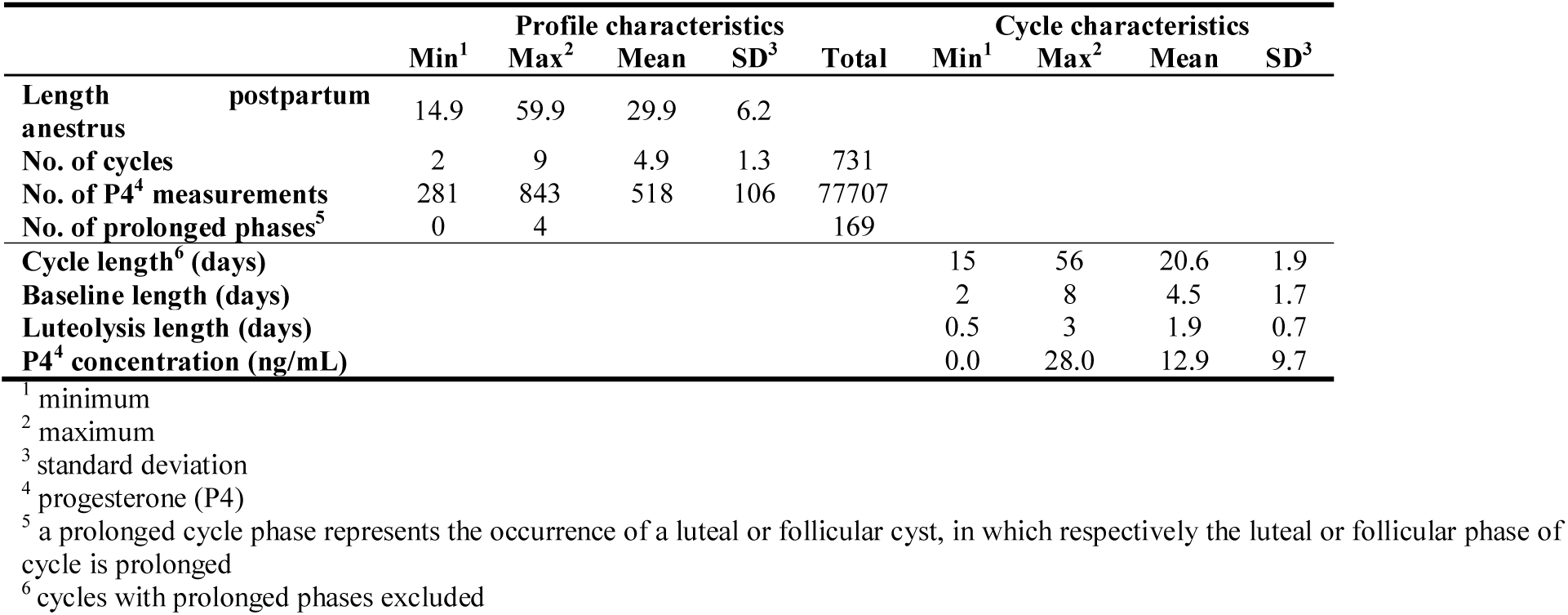
Progesterone (P4) profile and cycle characteristics of the simulated cows

**Figure 1.**
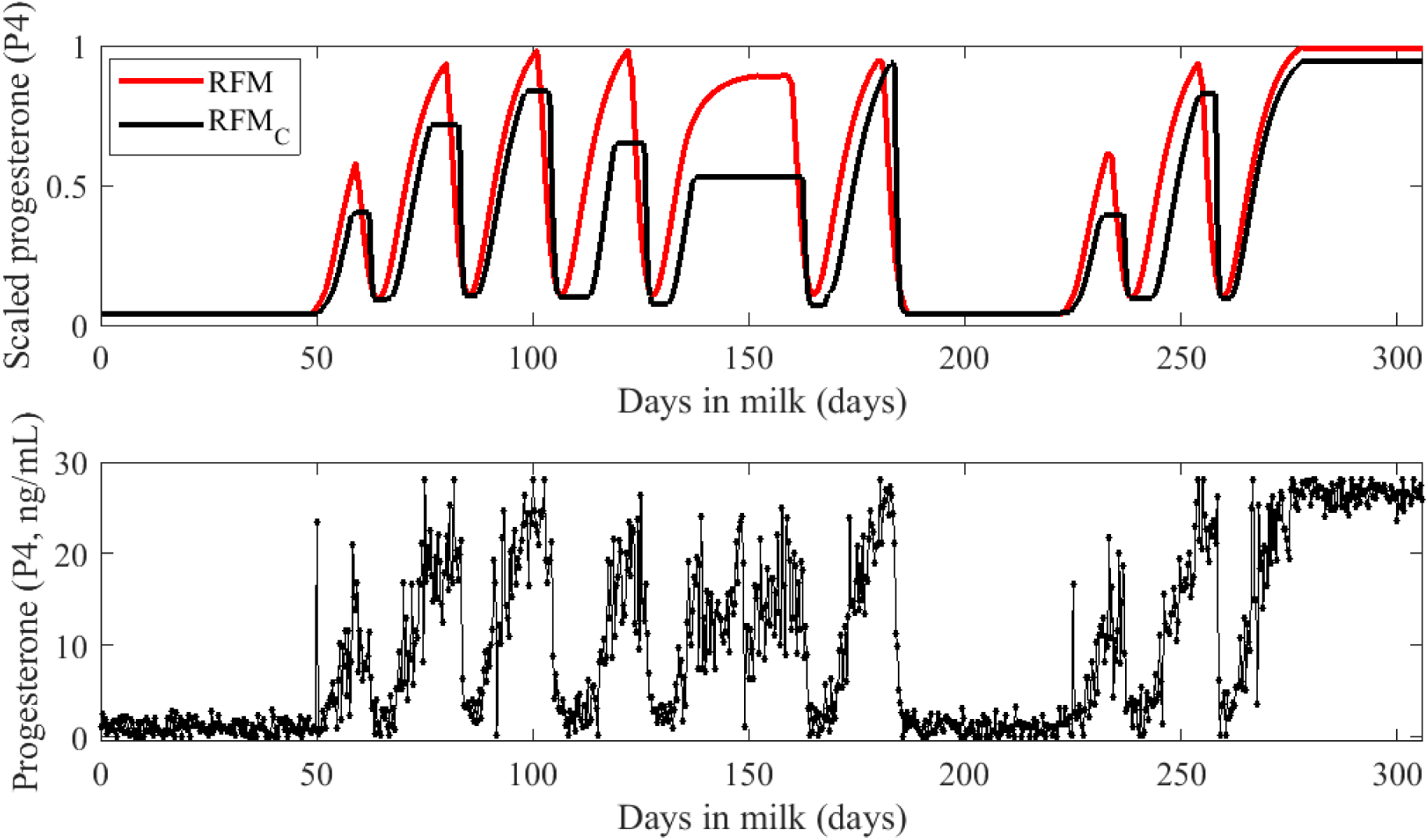
Upper panel: example of a simulated progesterone (P4) profile before (RFM, red) and after adding variability in profile characteristics (RFM_C_, black). Lower panel: resulting simulated P4 profile after descaling and adding measurement noise.

The outputs of the GARUNS/RFM simulations are scaled P4 profiles (i.e. representing reproduction performance over a lactation) of cows reflecting realistic fertility characteristics and with a measurement frequency of 1 sample per 2 hours (Martin et al., 2018). However, as these profiles are scaled, they do not represent the variability at cycle level (de-scaling), nor do they contain measurement noise associated with the P4 measurement in milk. Accordingly, two additional steps were required. The first step was to allow variability in the length and the relative P4 concentrations of the follicular and luteal phases. To this end, and to ensure maximal variability, the following properties of each cycle were adjusted according to a value randomly sampled from a uniform distribution, chosen according to the characteristics described by Meier et al. (2009a), Gorzecka et al. (2011) and Blavy et al. (2016):

1. The rate of decrease in P4 during luteolysis. In this step, the duration of the drop in P4 from maximum to minimum concentration was adjusted to last between 0.5 and 3 days (Gorzecka et al., 2011; Bruinjé et al., 2017). The uniform distribution used was thus U[0.5;3].
2. Adjustment of cycle height and shape. Meier and colleagues reported three different types of serum P4 profiles: ‘peaked’ profiles without a clear platform, ‘flat-top’ profiles with a distinguishable platform of constant high P4 production, and ‘structured’ profiles in which the P4 seems to rise in two phases (Meier et al., 2009b). To simulate these different types, the maximum P4 value in cycle was adjusted with a certain percentage varying from 40 to 100% (distribution U[0.4;1]) by either cutting off the data higher than this percentage or by multiplying the whole cycle with this percentage. The first procedure results in a ‘flat top’ shaped cycle, the second in a ‘peaked’ cycle. A structured shape was not considered, because in contrast to P4 in serum, this shape was not yet reported in milk.
3. The last step was to adjust the length of the baseline of each cycle from 3 to 8 days (Friggens and Chagunda, 2005; Blavy et al., 2016).

This procedure was repeated for each of the 731 cycles in the simulated dataset. An example of a (de-)scaled milk P4 profile is shown in black in the upper panel of Figure 1. For each simulated cycle, the reference moment of luteolysis was defined as the time (days in milk) at which the P4 level decreased below 70% of the difference between maximum and minimum P4 concentration within that cycle, further referred to as **REF_LUT_**. This moment of REF_LUT_ was chosen based on the assumption that the P4 concentration has to be sufficiently low before the dominant follicle can become LH sensitive. However occasionally, high-P4 estruses exist and thus full clearance of the P4 is not required (Friggens et al., 2008).

The second step entailed the addition of measurement noise corresponding to the on-line LFIA technique (resulting profile shown in lower panel of Figure 1). The characteristics of this measurement noise were defined from an available dataset containing 10,958 on-line measured P4 measurements collected at the Hooibeekhoeve in Geel, Belgium, which was described in (Adriaens et al., 2019). The P4 data in this LFIA dataset was smoothed using a second order Savitzky-Golay filter with a span of 7 measurements. Subsequently, the smoothed curve was subtracted from the data and the residuals were sorted based on their smoothed value, in which 28 classes of 1 ng/mL were created. For example, all measurements with a smoothed level between 0 and 1 ng/mL were assigned to the first class. Next, the standard deviation of the data in each class was calculated, and a second order polynomial was fitted to these standard deviation data, shown in Figure 2. This figure shows that the on-line measured P4 data is heteroscedastic, with less variability at the extremes than at intermediate values. The level of each simulated P4 measurement was obtained by multiplying the scaled simulated value with 28 ng/mL, while the second order polynomial fitted on the residuals was used to determine the standard deviation corresponding to this level. Next, a measurement was sampled from a normal distribution ~N(level, standard deviation). To introduce outliers in the dataset caused by e.g. a sampling error, one outlier on average each 8 days of measurements was sampled from a normal distribution with the maximum variance (i.e. a standard deviation of 5 ng/mL). This outlier represents e.g. milkings in which the sampling did not represent the whole milking (e.g. failed milkings or problems with the sampling unit). Accordingly, there was a chance of about 4% [= 1/ (8*3)] for each milking that the actual milk P4 value was replaced by an outlier sampled from a normal distribution with the maximum variance. A fixed milking interval of 8 hours (i.e. 3 samples per day) was chosen because the additional variability introduced by using realistic milking intervals (6 – 20 hours) would not have different results (own unpublished data). An overview of the cycle characteristics of the resulting dataset is given in the right part of Table 1.

**Figure 2.**
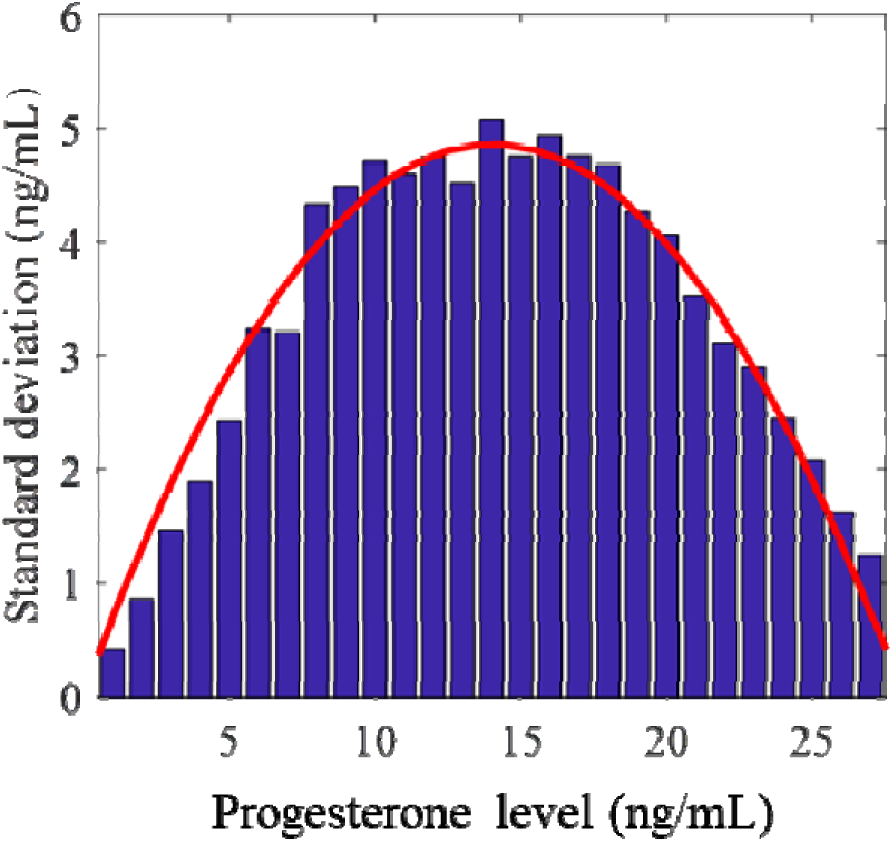
Distribution of measurement noise for when progesterone (P4) is determined via an on-farm lateral flow immune assay (LFIA) device. This device is optimized to discriminate between high and low P4 values and accordingly, has a low standard deviation at the extremes. The blue bars represent the standard deviation per smoothed progesterone concentration bin (e.g. 1-2 ng/mL), calculated from a real on-farm measured dataset and smoothed using a second order Savitzky-Golay filter with a span of 7 measurements. The red thick line is the fitted second order polynomial fitted on these standard deviations, and used to determine the simulated measurement noise for each progesterone measurement.

### Progesterone-based monitoring algorithm using synergistic control

The first algorithm tested in this paper, PMASC, consists of a mathematical model describing the luteal dynamics (Adriaens et al., 2017), and a statistical process control chart to detect luteolysis (Adriaens et al., 2018a). The mathematical model consists of two sigmoidal functions, a symmetrical Hill function to characterize the increase in P4 during luteal development, and a Gompertz function to describe the decrease during luteolysis. The control chart detects strong negative residuals from the predicted luteal P4 concentration, while taking the variability in P4 measurements into account. The decreasing function is only added after the detection of luteolysis, and can be used to calculate model-derived indicators that can be employed to inform the farmer on relevant actions to be taken (e.g. inseminations). It was shown before that the estimation of timing of luteolysis precedes the preovulatory LH surge by about 55 to 65 hours (Adriaens et al., 2018b). In this study, two outputs of PMASC were tested for their accuracy to relate to luteolysis. The first is the timing of the first measurement detected to be out-of-control (**OOC**), i.e. the exact time in hours that a first drop in P4 lower than expected is detected, followed by the confirmation of luteolysis in the two successive measurements. The second output ‘**TB85’** is an indicator calculated from the model as follows: the moment that the model describing the drop in P4 during luteolysis (i.e. the Gompertz function) decreases below 85% of the difference between maximum P4 model concentration minus the baseline; both calculated from the current P4 cycle characteristics (Adriaens et al., 2018b).

### Multi-process Kalman filter

To benchmark PMASC against the current state-of-the-art for on-line P4-based fertility monitoring, the simulated P4 values were smoothed using a MPKF as described in Friggens and Chagunda, (2005); Friggens et al., (2008); Løvendahl and Chagunda, (2010). More specifically, posterior mean estimates for the raw P4 values (i.e. the smoothed values) were calculated based on a ‘mixture’ of 4 local linear dynamic models. These local linear models represent the 4 possible ‘states’ in which the P4 time-series can be: steady state, encountering a slope or a level change, or an outlier. The mixture is calculated using the likelihood to be in a certain state taken from a predefined prior, and the one-step-back and two-steps-back posterior probabilities. This means that the reaction of the MPKF on a slope or level change increases with the extent of evidence for this state. For example, when the P4 values rapidly decrease from luteal to follicular concentrations, a first low measurement will be seen as very unlikely (‘outlier’), and the smoothed value will only decrease by a small amount. However, when the next sample is low again, there is more ‘evidence’ that this is a slope change, and an increased probability will be given to the ‘slope’ or ‘level’ change models, resulting in a larger decrease in the smoothed value.

The framework for the MPKF was set up based on the information provided in Smith and West, (1983), West and Harrison, (1997), Korsgaard and Løvendahl, (2002) and Friggens et al., (2007). The parameters were estimated based on the raw and smoothed P4 values of the same P4 dataset described before and in (Adriaens et al., 2019). The mean squared difference between our implementation of the MPKF and the smoothed values obtained from the on-line device was 0.094 ng/mL which was considered to be sufficiently low to compare results and derive relevant conclusions.

To provide a decision on when luteolysis has happened, the smoothed P4 values are combined with a fixed threshold to extract information from the time series. The threshold currently considered reasonable for estrus detection alerts based on milk P4 lies in between 4 and 6 ng/mL, and therefore in this study was taken 5 ng/mL (Friggens et al., 2008; Bruinjé et al., 2017). The MPKF+T algorithm gave an estrus alert when the smoothed value undercut the threshold value for the first time having previously exceeded this threshold value. To avoid multiple alerts within the same follicular period, a minimum time-interval of 5 days between two alerts was applied.

Once initiated and trained, this method does not need adjustments nor tuning to detect luteolysis. However, this user-friendly approach is at the cost of not including between-cow variation in responsiveness to P4. Moreover, the MPKF results in a time-lag between actual and detected luteolysis which is strongly dependent on luteolysis speed and the absolute P4 level measured (Friggens and Chagunda, 2005). A solution for this can be the implementation of a smart sampling scheme in which more samples are taken during the expected moment of luteolysis (e.g. from 16 days after the previous luteolysis on) and the calibration of the device to favor discrimination between high and low values.

### Detection performance

The simulated milk P4 dataset was analyzed using both algorithms (PMASC and MPKF+T) using 2 different sampling schemes. In the first sampling scheme, a milk P4 measurement was taken at each simulated milking (i.e. 3 times a day, sampling scheme ‘ALL’). For the second, only 1 sample per day (a i.e. 24h interval between samples) was provided to the algorithms (sampling scheme ‘1D’), which corresponded to a sample or data reduction of 66%. The latter mimics the effect of missing samples during luteolysis or a sampling scheme in which only 1 sample is taken per day during luteolysis to minimize the analysis costs. This is still a very high sampling rate, especially during the growth phase of the CL which is less of interest. Nevertheless, it gives an indication of what the performance can be with a sampling rate of only 1 sample per day in the period of expected luteolysis, while previously described studies sample once per milking in that period (Bruinjé et al., 2017). Next, the number and timing of simulated luteolyses was compared with the number of alerts given by each algorithm, and based hereon, the sensitivity, precision, and false negative rate (**FNR**) were calculated as follows (Eq. 1 to 3):

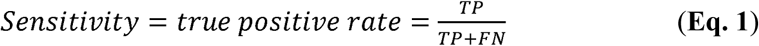

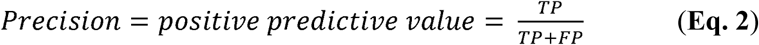

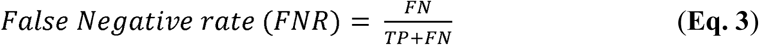

With **TP** being the true positives, i.e. the times PMASC or MPKF+T gave a luteolysis alert within a window of 2 days before (i.e. 6 samples for the ALL, and 2 samples for the 1D sampling scheme) to 4 days after (i.e. 12 samples for ALL and 4 samples for the 1D sampling scheme) a luteolysis was simulated (REF_LUT_). This window was chosen based on its practical relevance: detection later than 4 days would imply that ovulation had occurred by the time of detection, and the alert would be of no relevance for insemination (Roelofs, 2005; Adriaens et al., 2018b). False positives (**FP**) were alerts given by an algorithm which were not associated with a REF_LUT_. The false negatives (**FN**) were defined as cases where no alert was given at all or it was given later than 4 days after REF_LUT_. For this study, we chose not to include the specificity for a combination of reasons: (1) the specificity of the algorithms in this study is not fixed, unambiguously defined, but dependent on both the sampling rate and the window in which alerts are considered ‘true’ or ‘false’; (2) the specificity is not a measure where the farmer can do something with, as it is generally accepted that sensor systems should output as much as possible ‘farmers’ actions’, rather than raw data. Both concepts are further clarified in Hogeveen et al., (2010).

### Timing of the Alerts and Consistency and Robustness against Missing Samples

Besides sensitivity and accuracy measures, also the timing and variability in timing of the luteolysis alerts (i.e. P4 decrease) are of interest for an on-farm monitoring system. The first aspect, timing of the alert, is important because early alerts allow for correct planning of the insemination moment in order to achieve the highest chance of successful conception (Roelofs, 2005). The second aspect, variability in timing, is mainly of importance when a fixed insemination advice is coupled to the alerts. To this end, both the timing of the alerts and their variability compared to REF_LUT_ were evaluated for both sampling schemes. For PMASC, the first out-of-control measurement as well as the model-based indicator TB85 were included. The latter is calculated as the moment the decreasing Gompertz function reaches 85% of the difference between the between maximum and baseline P4 value of the model for that cycle, and thus takes the absolute milk P4 values of a cycle into account. This was shown previously to be the most consistent model-based decision criterion in relation to the LH surge (Adriaens et al., 2018b). Accordingly, 6 different groups were obtained, 3 for each sampling scheme: (1) the difference between REF_LUT_ and the first out-of-control measurement detected by PMASC, further referred to as OOC_ALL_ and OOC_1D_ respectively; (2) the difference between REF_LUT_ and TB85, referred to as TB85_ALL_ and TB85_1D_; and (3) the difference between REF_LUT_ and the timing of the milking that MPKF+T generated an alert, further indicated as MPKF+T_ALL_ and MPKF+T_1D_ for each of the sampling schemes, respectively.

Because of the unequal variances between the 6 different groups, we could not perform normal analysis of variance to compare their means. Therefore, the Wilcoxon Signed Rank Test was used to assess differences in median timing of the alerts, and the Brown-Forsythe test was used to investigate differences in variability. The latter tests the mean absolute difference from the median, which makes it robust for non-normality. However, because multiple comparison tests are not available for these statistical tests, it was decided to test each group against each other group in single pairwise comparisons. To avoid capitalization of chance due to multiple tests, a Bonferroni-correction on the significance level α=0.05 was applied by dividing it by the number of tests run, being 15 (each of the 6 groups compared to the 5 other groups). Differences were thus considered significant if the *p*-value was below 0.05/15 = 0.0033.

## RESULTS AND DISCUSSION

### Simulated data

The simulated dataset contained 150 milk P4 profiles and 731 individual cycles. Eighty-two cycles had a prolonged follicular phase (≥ 9 days), 53 cycles had a prolonged luteal phase (≥ 17 days) and 17 cycles had both. The average cycle length was 23.6 ± 7.6 days (mean ± SD) when all cycles were included, and 20.6 ± 1.9 days when cycles with prolonged phases were excluded. The average length of the follicular phase was 4.5 ± 1.7 days and the average duration of the P4 drop during luteolysis was 1.8 ± 0.7 days. The average baseline and maximum P4 concentrations were 2.1 ± 0.9 ng/mL and 18.8 ± 5.2 ng/mL, respectively. The characteristics of these profiles correspond to those obtained in a real on-farm setting and are in line with reported characteristics in the literature (Blavy et al., 2016). As such, the described methodology is a valuable way to generate large datasets while avoiding measurement costs and controlling both fertility and cycle characteristics while having a more precise reference for luteolysis. This study included in total 731 simulated luteolyses. Technically, simulating many more (e.g. a million) cycles was possible, but because of computational limitations, this current size was considered large enough number to show all possible difference between the groups (MPKF/PMASC and the 2 sampling schemes).

### Detection performance

In Table 2, the detection performance statistics for PMASC and MPKF+T are summarized for the different sampling schemes. When one measurement per milking was considered, PMASC and MPKF+T both had high sensitivities of 99.2% and 98.6%, respectively. For PMASC, also the precision of the alerts was high (95.3%), with only 4.7% of the 766 alerts being FP. With 23.0% FP alerts, the MPKF+T algorithm was more sensitive to variations in the data. More specifically, these false positive alerts can be classified as (1) outliers in the follicular phase (respectively 40 and 34% of the FP for 1D and ALL); (2) coincidentally successive measurements below the threshold, triggering the MPKF to low levels (respectively 43 and 49% of the FP for 1D and ALL); (3) cycles with intermediate maximal P4 concentration varying close to the threshold (17% of the FP for both 1D and ALL), which is often the case during e.g. luteal cysts (Yimer et al., 2018). To solve the first problem, an additional requirement in the MPKF+T model was that P4 had been above 15 ng/mL since the preceding estrus. The latter two situations are associated with the dependency of MPKF+T on the absolute measured P4 values, and are therefore not easily solved from a detection perspective. The MPKF+T of Friggens and Chagunda, (2005) gives an indication of the ‘goodness’ of the shape of the preceding cycle as information to aid the farmer in deciding whether to inseminate, which effectively flags these FPs (Friggens et al., 2008).

**Adriaens, Table 2.** Number of estruses detected by PMASC and MPKF+T, and the resulting sensitivity, precision and false negative rate (FNR) when the progesterone time series consisted of 3 samples per day (ALL) or 1 sample per day (‘1D’)

|  | PMASC<sup>8</sup> |  | MPKF+T<sup>9</sup> |  |
| --- | --- | --- | --- | --- |
|  | ALL<sup>6</sup> | 1D<sup>7</sup> | ALL | 1D |
| **Simulated** | **731** | **731** | **731** | **731** |
| **P<sup>1</sup>** | 761 | 742 | 937 | 808 |
| **TP<sup>2</sup>** | 725 | 723 | 721 | 704 |
| **FP<sup>3</sup>** | 36 | 19 | 216 | 104 |
| **FN<sup>4</sup>** | 6 | 8 | 10 | 27 |
| **Sensitivity (%)** | 99.2 | 98.9 | 98.6 | 96.3 |
| **Precision (%)** | 95.3 | 97.4 | 76.9 | 87.1 |
| **FNR<sup>5</sup> (%)** | 0.8 | 1.1 | 1.4 | 3.7 |

<sup>1</sup> P, positives. This is the total number of alerts for estrus
<sup>2</sup> TP, true positives. These are the alerts within the window of 2 days before and 4 days after a simulated luteolysis.
<sup>3</sup> false positives, FP. These are the alerts not associated with a simulated luteolysis
<sup>4</sup> False negatives, FN. These are simulated luteolyses not associated with an alert within the window of 2 days before and 4 days after a simulated luteolysis
<sup>5</sup> FNR, false negative rate
<sup>6</sup> ALL: dataset all contains all samples, corresponding to 1 measurement each 8 hours
<sup>7</sup> dataset 1D contains one sample per day
<sup>8</sup> PMASC: Progesterone Monitoring Algorithm using Synergistic Control
<sup>9</sup> MPKF+T: algorithm based on smoothed progesterone values using a multiprocess Kalman filter going below a fixed threshold of 5 ng/mL

Reducing the sampling frequency to one sample per day decreases the sensitivity for estrus detection of PMASC with 0.3% to 98.9%, and of MPKF+T with 2.2% to 96.4% (Table 2). The number of false detections were halved to 2.6% and 12.7%, respectively for PMASC and MPKF+T compared to the maximal sampling scheme, mainly because the number of outliers and coincidentally successive low values decreased. Correspondingly, the precision of MPKF+T increased with 10.3%. This shows the sensitivity of MPKF+T to the actual entered data and their absolute values. For PMASC, the FNR was very similar to that for the ‘ALL’ sampling scheme (1.1% and 0.8% respectively for ‘1D’ and ‘ALL’), which shows that the algorithm can work with less samples, as also presented in Adriaens et al., (2019). The MPKF+T seems to be more sensitive to this as the FNR increased from 1.4 to 3.6% when using less samples. The FN cases can be attributed to high-P4 estruses, in which samples close to the threshold were selected by coincidence. As a result, the smoothed values do not cross the threshold, or do not cross it in time (within a window of 4 days after simulated luteolysis). This phenomenon also shows the dependency of the MPKF+T on the actual absolute values. For example, when the P4 concentration drops significantly, e.g. from 25 ng/mL to 4.9 ng/mL, the MPKF tends to be conservative and will not drop to a value lower than the thresholds. Accordingly, at least 1 additional sample with a low P4 concentration has to be taken to trigger an alert. This time lag thereby ensures that real outliers do not immediately trigger an alert, which is important to guard the farmers’ trust. The advantage of PMASC is that its control limits are independent of the exact raw P4 values measured, and that the average value during the luteal phase is indirectly taken into account to monitor the drop during luteolysis. A similar observation was made by Friggens et al., (2008) when evaluating the luteolysis detection based on the model and algorithm described in Friggens and Chagunda, (2005). Although the authors started with a fixed threshold of 4 ng/mL, they had to implement another threshold of 6 ng/mL in order to detect high P4 estruses (Friggens et al., 2008). Because of the quite large measurement error for determining P4 in milk, it is not yet known whether the differences in P4 concentrations during estrus (and during the luteal phases) are due to inaccurate measurement and calibration methods, or due to real elevated P4 concentrations, e.g. due to increased P4 production by the adrenal cortex. When an improved P4 detection method becomes available, the inclusion of other information (e.g. health status, parity, fat content of the milk) might become possible and might improve the detection algorithms, but to date this is not yet available.

We did not test detection performance with other, more intelligent sampling schemes, because this was considered outside the scope of this study. First of all, other regular sampling schemes (e.g. 1 sample per 2 days) are irrelevant as detection would likely be too late for timely insemination, even if a follicular phase was detected. More specifically, PMASC was designed not just to ‘detect’, but also to ‘timely detect’ estruses in order to increase the chance for successful insemination.

### Timing of the Alerts: Variability and Robustness against Missing Samples

In Figure 3, the results of the timing of the alerts using all samples and one sample per day compared to REF_LUT_ are presented. The Wilcoxon signed rank test showed that not all medians are equal (*p* < 0.001), which can also be seen in Figure 3. Using all samples, the median difference between REF_LUT_ and the alerts generated by PMASC (TB85 and OOC) was close to 0, and their individual comparison test had a *p*-value larger than 0.44. More concretely, a median of zero for the OOC group means that the first indication of luteolysis (i.e. first sample out-of-control) is obtained almost simultaneously with the actual luteolysis REF_LUT_. This ‘early’ first indication allows to estimate the moment of luteolysis precisely, and has two large advantages: (1) it allows to organize the insemination at the best moment; and (2) it allows to wait for additional proof of real estrus or luteolysis in case of doubt, either by taking additional P4 samples, or by checking for external estrous symptoms. The latter can be of added value for example when farmers are skeptical about the reliability of new technologies. As expected, the most consistent insemination advice will be derived from a model-based indicator as can be calculated from PMASC, which allows determination of the moment of luteolysis independently of the sampling rate (when comparing 3 versus 1 sample per 24 hours). This paves the way to provide more consistent information to the farmer.

**Figure 3.**
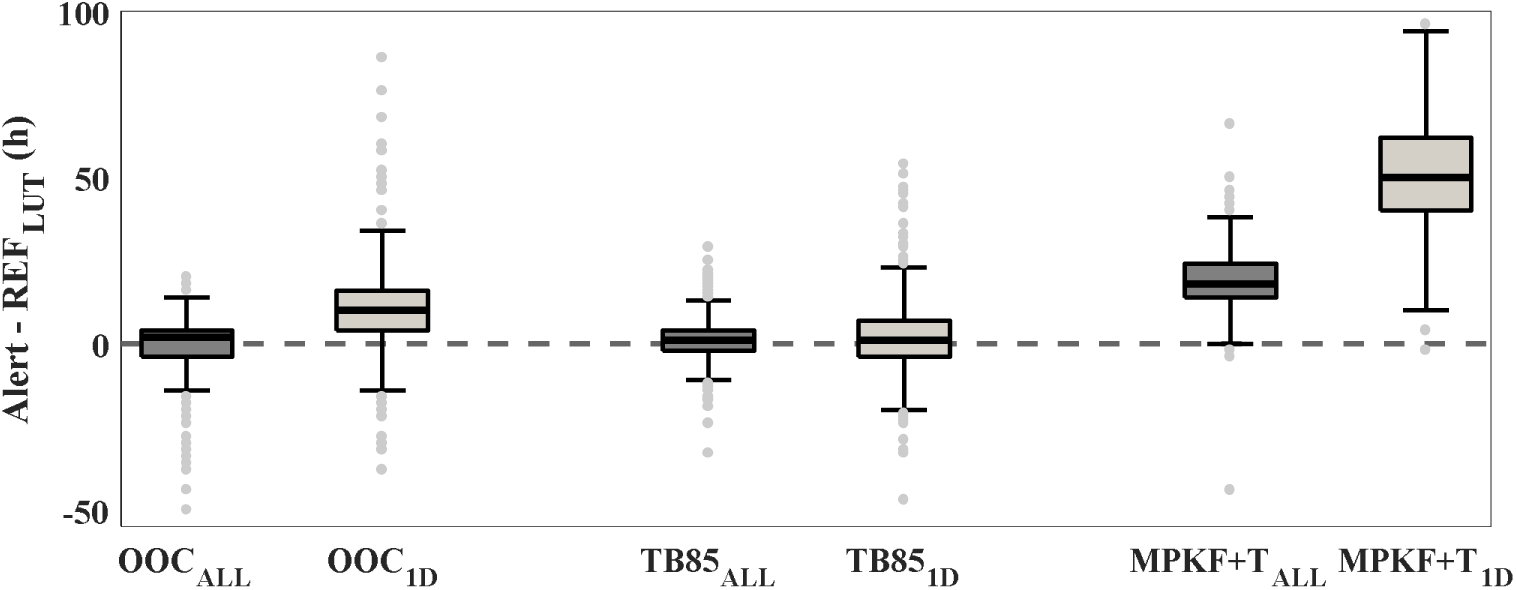
Boxplots for the differences between alerts given by the 2 algorithms included in this study (progesterone monitoring algorithm using synergistic control, PMASC and a multiprocess Kalman filter plus threshold, MPKF+T) and the simulated luteolysis (reference timing of luteolysis, REF_LUT_). A difference of zero means that luteolysis is detected on the moment it is simulated, which is seen for the first out-of-control sample of PMASC using all samples (first out-of-control, milk progesterone simulated 3 times a day, OOC_ALL_) and for alerts based on the model-derived indicator TB85, even when samples during luteolysis are missed (TB85_ALL_ and TB85_1D_). ‘1D’ hereby means that only 1 sample per day was taken, also during the period in which luteolysis occurred.

For the OOC_1D_ group (first out-of-control measurement detected by PMASC for the one-sample-per day scheme), the first indication of estrus is obtained on average 10 hours later (significantly different from all other groups, *p* < 0.001). However, using TB85_1D_, (i.e. the model-based indicator using the maximum and minimum model P4 value) estimating the actual moment of luteolysis in a consistent way close to REF_LUT_ remains possible. The alerts of MPFK+T_ALL_ and MPKF+T_1D_ came respectively on average 18 and 48 hours after REF_LUT_, which also were significantly different from all other medians (*p* < 0.001). Accordingly, the MPKF+T algorithm has a time lag in detection of about 2 milkings when no samples are missed during luteolysis, while this lag increases to approximately 2 days when only one sample per day is taken. As a result, it becomes difficult to provide the farmer with a reliable estimation of optimal insemination time, which is not only dependent on the number of samples missed, but also on how fast P4 decreased (and thus, the MPKF reacts), the moment of luteolysis compared to the timing of the milkings and milking intervals.

Table 3 shows that the Brown-Forsythe test for differences in variability in the interval alert to REF_LUT_ was highly significant. The variability for TB85_ALL_ was the smallest, with a standard deviation respectively 35.3% and 15.8% smaller than OOC_ALL_ and MPKF+T_ALL_. The OOC_ALL_ and MPKF+T_ALL_ groups had an equal variability (*p*-value = 0.017 > 0.0033). Furthermore, individual comparisons pointed out that the variability does not differ between the OOC and TB85 group when one sample per day was considered (*p-*value = 0.047 > 0.0033). In contrast, we see that missing samples during luteolysis have a larger effect on the consistency of alerts given by the MPKF+T algorithm than on those obtained from OOC and TB85, resulting in a significantly larger variability than all other groups (*p*-value < 0.0033). We can therefore conclude that OOC and TB85 are less sensitive to missing samples both in terms of median timing of alerts and in terms of variability, making it a more robust algorithm for missing samples during luteolysis compared to MPKF+T.

**Adriaens, Table 3.**
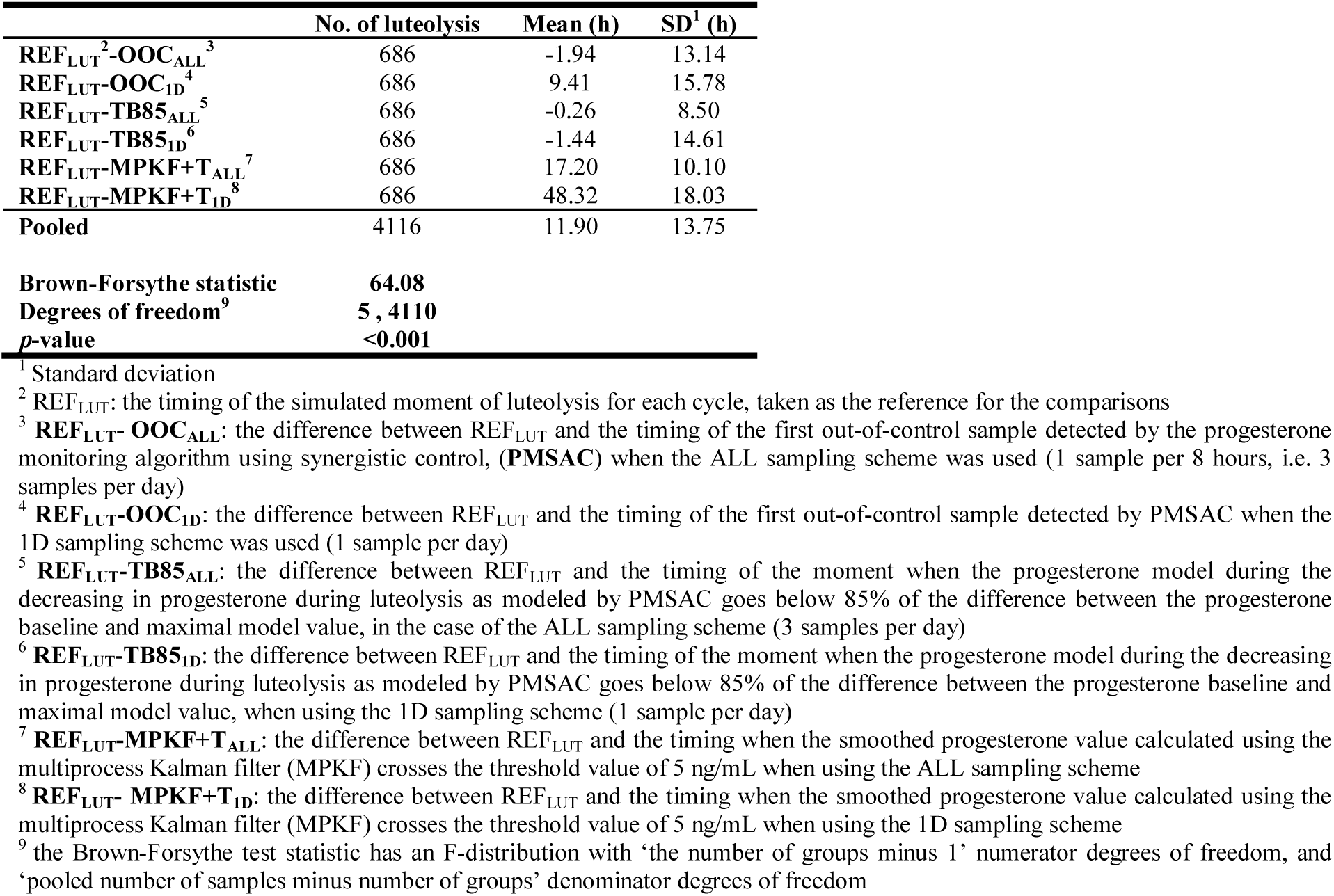
Overview of mean, standard deviation and the outcome of the Brown-Forsythe test for equal absolute deviations of the median for all comparisons for the different algorithms (the progesterone monitoring algorithm using synergistic control, PMSAC and the multiprocess Kalman filter + a fixed threshold, MPKF+T) and sampling schemes (ALL: three times a day sampling vs. 1D: once a day sampling).

Part of the variability within the groups is caused by the timing of the milkings relative to the P4 profiles. For example, in this study a sample could have been taken in a window of 0 to 8 hours after REF_LUT_. Accordingly, the effect of the timing of milking relative to REF_LUT_ seems to overrule the effect of the algorithm for the scheme in which milk P4 is measured every 8 hours. Although taking only one sample per day might not seem a ‘random’ way to mimic missing samples during luteolysis, the variability in luteolysis length and the independency of the P4 profiles to the simulated time of REF_LUT_ ensures that the timing of missed samples compared to luteolysis was variable. In a real on-farm setting, it is more probable that not in all cows samples are skipped, which would make the variability in alerts compared to real luteolysis for OOC and MPKF+T even larger (see also Adriaens et al., 2019). Furthermore, based on the results of this study, we can assume that the TB85 indicator remains consistent in its estimation of the timing luteolysis, independent of the sampling scheme or interval, which supports its use for monitoring purposes. A robust indicator is important when a fixed rule is used for decision making.

## CONCLUSIONS

The detection performance of PMASC and the MPKF+T in terms of sensitivity, precision, false negative rate, false detection rate, robustness for sampling frequency and missing samples during luteolysis were studied on a simulated dataset of 150 highly variable P4 profiles containing 731 luteolyses preceding estrus. Both PMASC and MPKF+T had a high luteolysis detection rate, but MPKF+T was more sensitive to the absolute values of the P4 data, shown by its higher false detection rate. This illustrates the value and limitations of both algorithms for on-line fertility monitoring. Using a PMASC-based model indicator taking into account the luteal and follicular P4 concentrations, a more robust estimation of the timing of luteolysis was obtained, which was less sensitive to missing values compared to the current state-of-the-art MPKF+T both in terms of detection rate and variability. Accordingly, PMASC has shown its potential to improve consistency and robustness of progesterone-based, cost-effective detection of luteolysis on farm.

## ACKNOWLEDGEMENTS

This work was supported by the Institute for the Promotion of Innovation through Science and Technology in Flanders, Belgium (IWT) [IWT-LA project 110770]. Ines Adriaens and Ben Aernouts are supported by the Fund for Scientific Research (FWO) Flanders, respectively grant number 11ZG916N and 12K3916N. Ines Adriaens obtained additional funding to perform a research stay at INRA, grant V410318N. The RFM model was developed within the ‘Project Pluridisciplinary study for a RObust and sustainabLe Improvement of Fertility In Cows’, PROLIFIC, EU grant no. FP7.KBBE.2012.6, grant agreement no. 311776.

## APPENDIX

**Figure A1.**
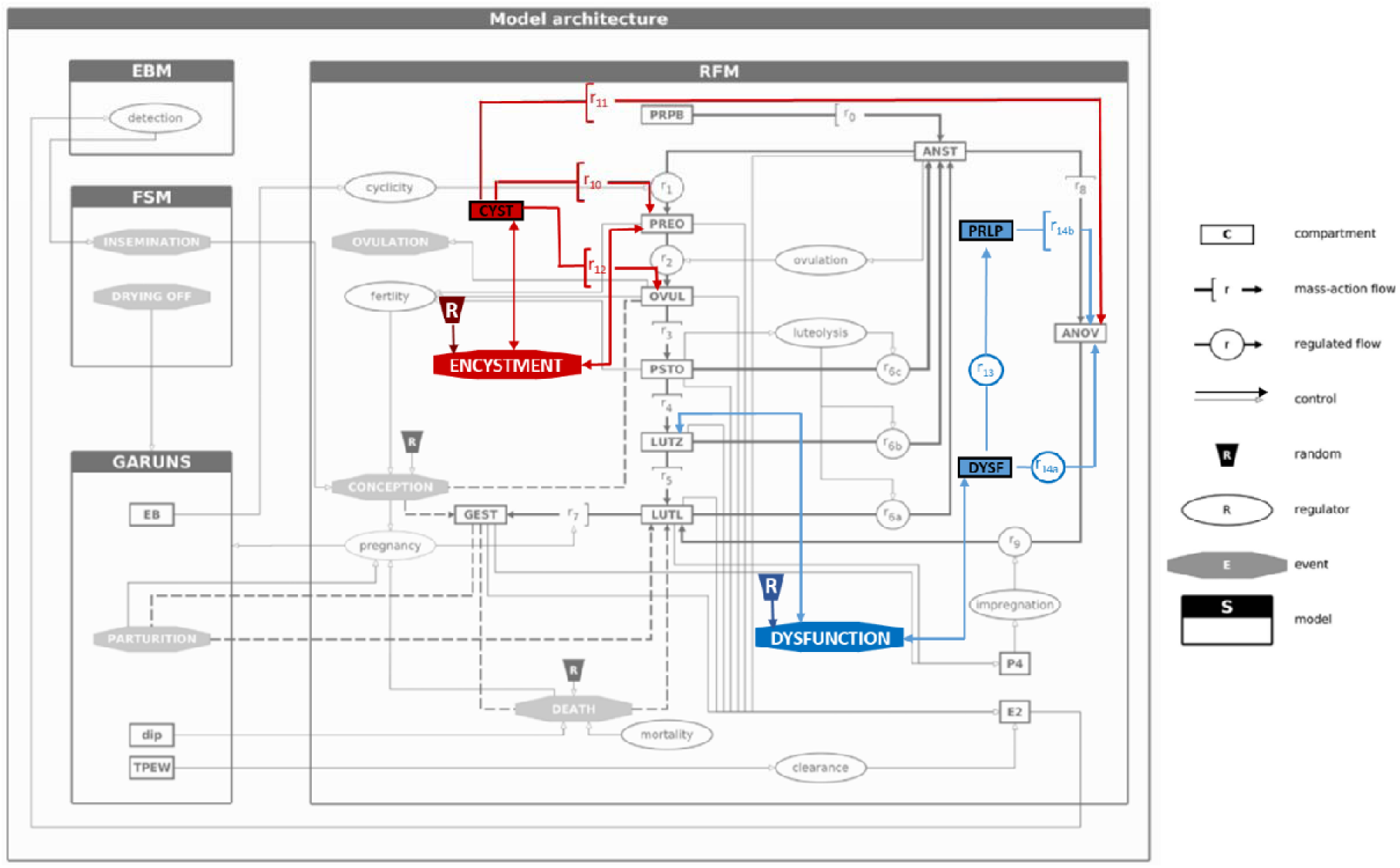
Model structure of the Reproduction Function Model and the adaptations made to allow for simulation of interrupted cyclicity by prolonged luteal (DYSF and PRLP, indicated in blue) and follicular phases (CYST, indicated in red), adapted from Martin et al., (2018).

**Adriaens, Table A1.** Parameters for the reproduction function model (RFM) model as described in Martin et al., (2018). The variability in genetic scaling parameters allows to alter individual performance of cows at the level of GARUNS. To allow for additional variability in fertility performance of the simulated cows, two sets of parameters (set 1 and set 2) were entered in the model. Using this table, new progesterone profiles can be simulated in a similar way as done in this study.

| Parameter name RFM<sup>1</sup> | Set 1 | Set 2 | Effect |
| --- | --- | --- | --- |
| **h_CYCL**<sup>2</sup> | **-7.0** | **-15.0** | Adjust postpartum anestrus length |
| **Risk_CYST**<sup>3</sup> | **1.6** | **2.0** | Incidence of follicular cysts |
| **Risk_DYSF**<sup>4</sup> | **1.6** | **2.0** | Incidence of luteal dysfunction |
| **EMO**<sup>5</sup> | **2.1** | **2.5** | Incidence early embryonic mortality (conception failure) |
| GSP_cv<sup>6</sup> | 0.05 | 0.05 | Variability between cows & characteristics (genetic scaling parameters) |
| RFM_h_impregnation[4]<sup>7</sup> | 0.10 | 0.10 | Ensure restart of cyclicity after R_CYST |
| RFM_k09<sup>8</sup> | 0.10 | 0.10 | Ensure restart of cyclicity after R_CYST |
| RFM_k10<sup>9</sup> | 0.12 | 0.12 | Duration follicular cyst / ovarian anestrus |
| RFM_k11<sup>10</sup> | 0.12 | 0.12 | Duration follicular cyst / ovarian anestrus |
| RFM_k13<sup>11</sup> | 0.3 | 0.3 | Duration luteal dysfunction |
| RFM_EMO[2]<sup>12</sup> | 4.6 | 4.6 | Avoid occurrence late embryonic mortality |
| RFM_EMO[3]<sup>13</sup> | 8.0 | 8.0 | Avoid occurrence fetal death |

<sup>1</sup> RFM: reproduction function model, the model used for simulating progesterone profiles in this study
<sup>2</sup> Parameter determining the length of the postpartum anestrus phase. One of the 4 parameters varied for both simulations (set 1 and set 2)
<sup>3</sup> Parameter determining the incidence of follicular cysts. One of the 4 parameters varied for both simulations (set 1 and set 2)
<sup>4</sup> Parameter determining the incidence of luteal dysfunction. One of the 4 parameters varied for both simulations (set 1 and set 2)
<sup>5</sup> Parameter determining the incidence of early embryonic mortality. One of the 4 parameters varied for both simulations (set 1 and set 2)
<sup>6</sup> GSP\_cv: coefficient of variation of the genetic scaling parameters. This parameter determines how much difference there is between simulated cows, and is indicative for the absolute values of some of the other outcomes (milk production, body weight) of the GARUNS model
<sup>7</sup> One of the 2 parameters that ensure cyclicity restarts in the model after an interruption due to a prolonged follicular period (low progesterone)
<sup>8</sup> One of the 2 parameters that ensure cyclicity restarts in the model after an interruption due to a prolonged follicular period (low progesterone)
<sup>9</sup> One of the 2 parameters of the RFM model influencing the duration of follicular problems, reflected in prolonged follicular phases of the progesterone cycle (low progesterone)
<sup>10</sup> One of the 2 parameters of the RFM model influencing the duration of follicular problems, reflected in prolonged follicular phases of the progesterone cycle (low progesterone)
<sup>11</sup> Parameter influencing the duration of luteal dysfunction, reflected in prolonged luteal phases of the progesterone cycle (high progesterone)
<sup>12</sup> Parameter determining when and how often late embryonic mortality occurs
<sup>13</sup> Parameter determining when and how often fetal death occurs

## REFERENCES

Adriaens, I., T. Huybrechts, K. Geerinckx, D. Daems, J. Lammertyn, B. De Ketelaere, W. Saeys, and B. Aernouts. 2017. Mathematical characterization of the milk progesterone profile as a leg up to individualised monitoring of reproduction status in dairy cows. Theriogenology 103:44–51. doi: https://doi.org/10.1016/j.theriogenology.2017.07.040.

Adriaens, I., W. Saeys, K. Geerinckx, B. De Ketelaere, and B. Aernouts. 2019. Short communication: Validation of a novel milk progesterone based tool to monitor luteolysis in dairy cows. Performance on cost-effective, on-farm measured data. J. Dairy Sci. [accepted]. doi: https://doi.org/10.1101/526061.

Adriaens, I., W. Saeys, T. Huybrechts, C. Lamberigts, K. Geerinckx, J. Leroy, B. De Ketelaere, and B. Aernouts. 2018a. A novel system for online fertility monitoring based on milk progesterone. J. Dairy Sci. 101:1–14. doi: 10.3168/jds.2017-13827.

Adriaens, I., W. Saeys, C. Lamberigts, M. Berth, K. Geerinckx, J. Leroy, B. De Ketelaere, and B. Aernouts. 2018b. Short communication: Sensitivity of estrus alerts and relationship with timing of the luteinizing hormone surge. J. Dairy Sci. 102:1–5. doi: https://doi.org/10.3168/jds.2018-15514.

Blavy, P., M. Derks, O. Martin, J.K. Hoglund, and N.C. Friggens. 2016. Overview of progesterone profiles in dairy cows. Theriogenology 86:1061–1071. doi: https://doi.org/10.1016/j.theriogenology.2016.03.037.

Braw-Tal, R., S. Pen, and Z. Roth. 2009. Ovarian cysts in high-yielding dairy cows. Theriogenology 72:690–698. doi: 10.1016/j.theriogenology.2009.04.027.

Bruinjé, T.C., M. Gobikrushanth, M.G. Colazo, and D.J. Ambrose. 2017. Dynamics of pre- and post-insemination progesterone profiles and insemination outcomes determined by an in-line milk analysis system in primiparous and multiparous Canadian Holstein cows. Theriogenology 102:147–153. doi: https://doi.org/10.1016/j.theriogenology.2017.05.024.20

Friggens, N.C., M. Bjerring, C. Ridder, S. Højsgaard, and T. Larsen. 2008. Improved detection of reproductive status in dairy cows using milk progesterone measurements. Reprod. Domest. Anim. 43:113–121.

Friggens, N.C., and M.G.G. Chagunda. 2005. Prediction of the reproductive status of cattle on the basis of milk progesterone measures: model description. Theriogenology 64:155–90. doi: 10.1016/j.theriogenology.2004.11.014.

Friggens, N.C., K.L. Ingvartsen, I.R. Korsgaard, T. Larsen, P. Løvendahl, C. Ridder, and N.I. Nielsen. 2007. System and a method for observing and predicting a physiological state of an animal. US Pat. No. US7302349 B2.

Gaillard, C., O. Martin, P. Blavy, N.C. Friggens, J. Sehested, and H.N. Phuong. 2016. Prediction of the lifetime productive and reproductive performance of Holstein cows managed for different lactation durations, using a model of lifetime nutrient partitioning. J. Dairy Sci. 99:9126–9135. doi: 10.3168/jds.2016-11051.

Gorzecka, J., M.C. Codrea, N.C. Friggens, and H. Callesen. 2011. Progesterone profiles around the time of insemination do not show clear differences between of pregnant and not pregnant dairy cows. Anim. Reprod. Sci. 123:14–22. doi: 10.1016/j.anireprosci.2010.11.001.

Hogeveen, H., C. Kamphuis, W. Steeneveld, and H. Mollenhorst. 2010. Sensors and Clinical Mastitis—The Quest for the Perfect Alert. Sensors 10:7991–8009. doi: 10.3390/s100907991.

Jeengar, K., V. Chaudhary, A. Kumar, S. Raiya, M. Gaur, and G.N. Purohit. 2014. Ovarian cysts in dairy cows: old and new concepts for definition, diagnosis and therapy. Anim. Reprod. 11:63–73.

Korsgaard, I.R., and P. Løvendahl. 2002. An introduction to multiprocess class II mixture models. Pages 185-188 in Proceedings of the 7th World Congress on genetics applied to livestock production. Département de Génétique Animale INRA, Castanet-Tolosan.

Løvendahl, P., and M.G.G. Chagunda. 2010. On the use of physical activity monitoring for estrus detection in dairy cows. J. Dairy Sci. 93:249–259. doi: 10.3168/jds.2008-1721.

Martin, O., P. Blavy, M. Derks, N. Friggens, and F. Blanc. 2018. Coupling a reproductive function model to a productive function model to simulate lifetime performance in dairy cows. Animal 1:1–10. doi: 10.1017/S1751731118001830.

Martin, O., N.C. Friggens, J. Dupont, P. Salvetti, S. Freret, C. Rame, S. Elis, J. Gatien, C. Disenhaus, and F. Blanc. 2013. Data-derived reference profiles with corepresentation of progesterone, estradiol, LH, and FSH dynamics during the bovine estrous cycle. Theriogenology 79:331–343. doi: https://doi.org/10.1016/j.theriogenology.2012.09.025.

Martin, O., and D. Sauvant. 2010. A teleonomic model describing performance (body, milk and intake) during growth and over repeated reproductive cycles throughout the lifespan of dairy cattle. 1. Trajectories of life function priorities and genetic scaling. Animal 4:2030–2047. doi: 10.1017/S1751731110001357.

Meier, S., J. Roche, E. Kolver, G. Verkerk, and R. Boston. 2009a. Comparing subpopulations of plasma progesterone using cluster analyses. J. Dairy Sci. 92:1460–1468. doi: 10.3168/jds.2008-1464.

Meier, S., J.R. Roche, E.S. Kolver, and R.C. Boston. 2009b. A compartmental model describing changes in progesterone concentrations during the oestrous cycle. J. Dairy Res. 76:249–56.

Peter, A.T., H. Levine, M. Drost, and D.R. Bergfelt. 2009. Compilation of classical and contemporary terminology used to describe morphological aspects of ovarian dynamics in cattle. Theriogenology 71:1343–1357. doi: 10.1016/j.theriogenology.2008.12.026.

Ranasinghe, R.M.S.B.K., T. Nakao, K. Yamada, K. Koike, A. Hayashi, and C.M.B. Dematawewa. 2011. Characteristics of prolonged luteal phase identified by milk progesterone concentrations and its effects on reproductive performance in Holstein cows.. J. Dairy Sci. 94:116–127.

Roelofs, J.B. 2005. When to inseminate the cow? Insemination, ovulation and fertilization in dairy cattle. PhD thesis, Wageningen Institute of Animal Sciences, Wageningen, the Netherlands,. 22

Rosenberg, L. 2010. Cystic ovaries in dairy cattle. PhD thesis, Dairy Science Department, California Polytechnic State University, San Luis Obispo,.

Smith, A., and M. West. 1983. Monitoring renal transplants: an application of the multiprocess Kalman filter. Biometrics 39:867–878.

West, M., and J. Harrison. 1997. Bayesian Forecasting and Dynamic Models. 2nd ed. Springer US, New York.

Yimer, N., A. Haron, and R. Yusoff. 2018. Determination of Ovarian Cysts in Cattle with Poor Reproductive Performance Using Ultrasound and Plasma Progesterone Profile. Vet. Med. 3:1–9. doi: 10.1017/dmp.2016.67.

